# Spatio-Temporal Correlates of Gene Expression and Cortical Morphology across Life Course and Aging

**DOI:** 10.1101/2020.07.16.207753

**Authors:** Anqi Qiu, Han Zhang, Brian K. Kennedy, Annie Lee

## Abstract

Evidence from independent neuroimaging and genetic studies supports the concept that brain aging mirrors development. However, it is unclear whether mechanisms linking brain development and aging provide new insights to delay aging and potentially reverse it. This study determined biological mechanisms and phenotypic traits underpinning brain alterations across the life course and in aging by examining spatio-temporal correlations between gene expression and cortical volumes (n=3391) derived from the life course dataset (3-82 years) and the aging dataset (55-82 years). We revealed that a large proportion of genes whose expression was associated with cortical volume across the life course were in astrocytes. These genes, which showed up-regulation during development and down-regulation during aging, contributed to fundamental homeostatic functions of astrocytes crucial, in turn, for neuronal functions. Included among these genes were those encoding components of cAMP and Ras signal pathways, as well as retrograde endocannabinoid signaling. Genes associated with cortical volumes in the aging dataset were also enriched for the sphingolipid signaling pathway, renin-angiotensin system (RAS), proteasome, and TGF-beta signaling pathway, which is linked to the senescence-associated secretory phenotype. Neuroticism, drinking, and smoking were the common phenotypic traits in the life course and aging, while memory was the unique phenotype associated with aging. These findings provide biological mechanisms and phenotypic traits mirroring development and aging as well as unique to aging.

## Introduction

The Developmental Theory of Aging (DTA) (1–4) has recently received great attention. It posits that brain aging may be a reverse process of brain development. Indeed, neurogenesis, myelination, and synaptic pruning in early life are mirrored by neuronal loss, demyelination, and synaptic loss in aging. Nevertheless, strategies for reversal of brain aging have not been successfully developed based on this or any other theory. Understanding the mechanisms behind the DTA may lead to such strategies.

Evidence from independent neuroimaging and genetic studies suggests a possible link between brain development and aging, supporting the idea of DTA. Neuroimaging studies suggested that brain volume loss in normal aging mirrors brain volume gain in development (3, 4). For instance, brain regions that develop later degenerate earlier (3, 4). The transcriptome has been meticulously explored in a large collection of human brain specimens spanning from infant development to old age (1, 2). The majority of gene expression changes in aging occur in the opposite direction of gene expression changes in postnatal development (5). In other words, many genes that show up (or down)-regulation during development also show down (or up)-regulation during aging. This reversal of gene expression across the life course is suggested to be a genomic reflection that bridges developmental and aging processes (2). Genes that are up-regulated during aging to the levels observed in development may signify a response to oxidative DNA damage (6, 7). Similarly, genes with an up-down expression pattern across the life course contribute to biological processes related to neuronal and synaptic functions (5). The reversal may be possibly associated with aging-related accumulation of stochastic effects that lead to reduced maintenance of mature, differentiated cells and provide a possible link to the brain and cognitive aging (8). These biological processes have been highlighted as potential factors relevant to senescent processes in aging (9). Nevertheless, senescence during brain postnatal development remains barely understood (10).

This study explored several questions: 1) do genes with reverse expression across the life course belong to specific cell types? 2) do they contribute to senescent processes common in brain development and aging? 3) are there any processes only in brain aging? 4) what phenotypic traits associated with these genes can be targets for early prevention of aging? To answer these questions, we integrated neuroimaging and transcriptome data, examining the similarity of cortical morphology and gene expression throughout the brain and across age. We further employed a series of bioinformatics tools to identify brain-tissue, cell-specific genes and their contributions to biological pathways and phenotypic traits common in development and aging (life course: age range from 3 to 82 years) or unique to aging (55 years above). We expected that integrating neuroimaging and transcriptome data would provide new insights on DTA. More specifically, the genes strongly correlated with the spatio-temporal pattern of cortical morphology can identify senescent pathways and phenotypic traits linking development and aging. Together, our findings provide potential targets for future investigation that may aid the development of interventions to prevent the age-related decline in brain function.

## Results

In this study, we used the term “life course” to represent the age range from 3 to 82 years and the term “aging” to denote the age range from 55 years and above.

### The transcriptome of the cortex across the life course

We first examined the transcriptional architecture throughout the brain and across the life course (age from 3 to 82 years). We utilized the transcriptome dataset published by Kang and colleagues(11). This included expression data of 17,565 genes from 26 donors aged 3 to 82 years, sampled from 11 cortical regions, including the orbitofrontal cortex, ventrolateral frontal cortex, dorsolateral frontal cortex, medial frontal cortex, precentral and postcentral gyri, transverse temporal gyrus, superior and inferior temporal gyri, supramarginal gyrus, and calcarine (**Supplementary Table 1**).

A quadratic regression model was used to examine age effects on gene expression across the life course. We identified genes whose temporal pattern across the life course is either down-up or up-down for all the cortical regions. For instance, **Figures 1A** illustrates that the MAPK1 gene was up-regulated during postnatal development and then down-regulated in aging, displaying an up-down (or reversal) pattern across the life course and all 11 cortical regions. 710 genes were identified with the up-down or down-up expression pattern across the life course and all the 11 cortical regions.

**Figure 1.**
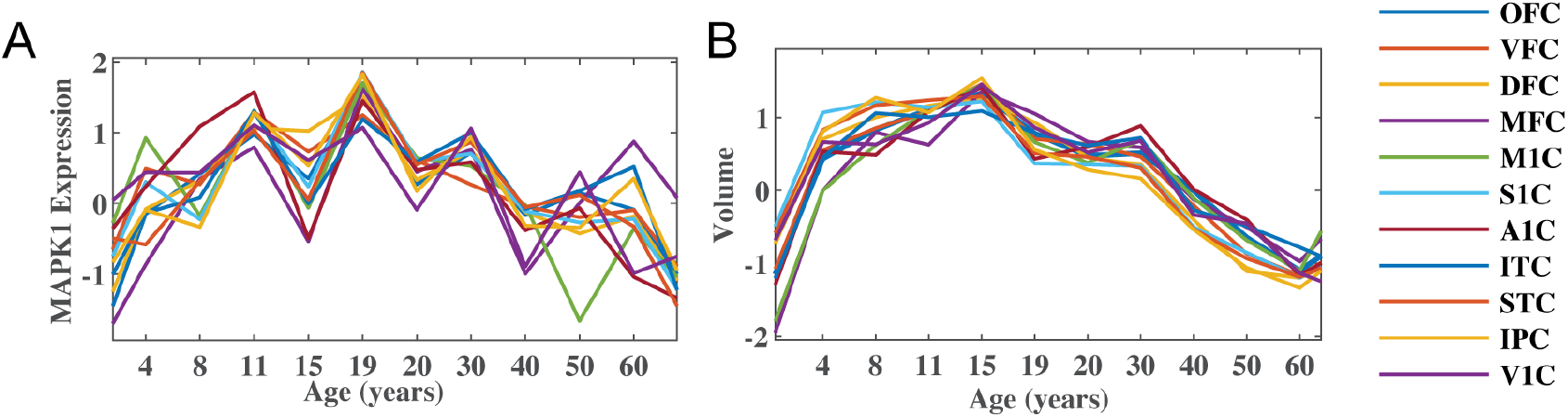
Gene expression and brain volumes as a function of age. A. MAPK1 expression in the cortex; B. cortical volumes. Abbreviations: OFC, orbitofrontal cortex; VFC, ventrolateral prefrontal cortex; DFC, dorsolateral prefrontal cortex; MFC, medial frontal cortex; M1C, motor cortex; S1C, somatosensory cortex; A1C, transverse temporal cortex; ITC, inferior temporal cortex; STC, superior temporal cortex; IPC, inferior parietal cortex; V1C, calcarine.

### Morphological pattern of the brain across the life course

The cortical volumes were estimated from the T_1_-weighted MRI of 2765 subjects aged 3 to 82-years-old using FreeSurfer (**Section A in the Supplementary Material**). **Supplementary Table 5** summarizes the demographics of these subjects. **Supplementary Figure 1** shows scatter plots of the 11 cortical volumes in relationship with age. Quadratic linear regression analysis revealed all 11 cortical volumes as a function of age^2^, suggesting a significant inverted-U relationship with age across the life course in all these cortical regions (corrected *p*<0.001).

### Spatio-temporal correlation between gene expression and brain morphology across the life course

We aimed to identify genes whose spatio-temporal profiles are similar to spatio-temporal patterns of the cortical morphology across the life course. We then investigated the contribution of these genes to biological pathways that may be relevant to brain development and aging. For this, we employed spatio-temporal correlation analysis to examine the relationships between gene expression and brain volumes within the cortex across the life course.

Among the 710 genes with up-down or down-up gene expression profiles in the cortex, 574 genes showed significant spatio-temporal correlation with the cortical volumes (corrected *p*<0.05, see an example in **Figure 1**). TISSUES 2.0(12) was used to compute a confidence score and to identify genes that were preferentially expressed in the brain compared to other tissues. The confidence score reflects the likelihood of a true association between genes and a given tissue type. A confidence score of 4.5 or higher was used to define brain-dominant genes in this study (12). Among 574 genes, 293 genes were brain-dominant (e.g., BDNF, CDC42, MAPK1, MAPK9, etc) and 261 of them had the up-down expression pattern across the life course, similar to that of MAPK1 in **Figure 1A**. **Figure 2A** shows the distribution of cell types of these brain-dominant genes, including 240 astrocytes (226 in the up-down pattern), 14 neurons (11 in the up-down pattern), 20 oligodendrocytes (13 in the up-down pattern), 14 microglia (7 in the up-down pattern), and 5 unclassified cells. Fisher’s test revealed that 260 astrocyte genes were overrepresented among 293 genes when the brain genes studied here were used as reference (p<0.001). **Table S2** lists the genes in each cell type. The genes in neurons, oligodendrocytes, and microglia were not enriched for any biological pathway. The astrocytes (n=240) were found to be enriched for biological pathways (**Table S3**) that can be classified into 4 categories, including retrograde endocannabinoid, oxytocin, Ras, and cAMP signaling pathways (all *p*<0.001, see **Figure 3A** and **3B**). **Figure 3C** shows the genes contributing to individual biological pathways, annotates the genes with the up-down pattern across the life course in red and the genes with the down-up pattern in blue.

**Figure 2.**
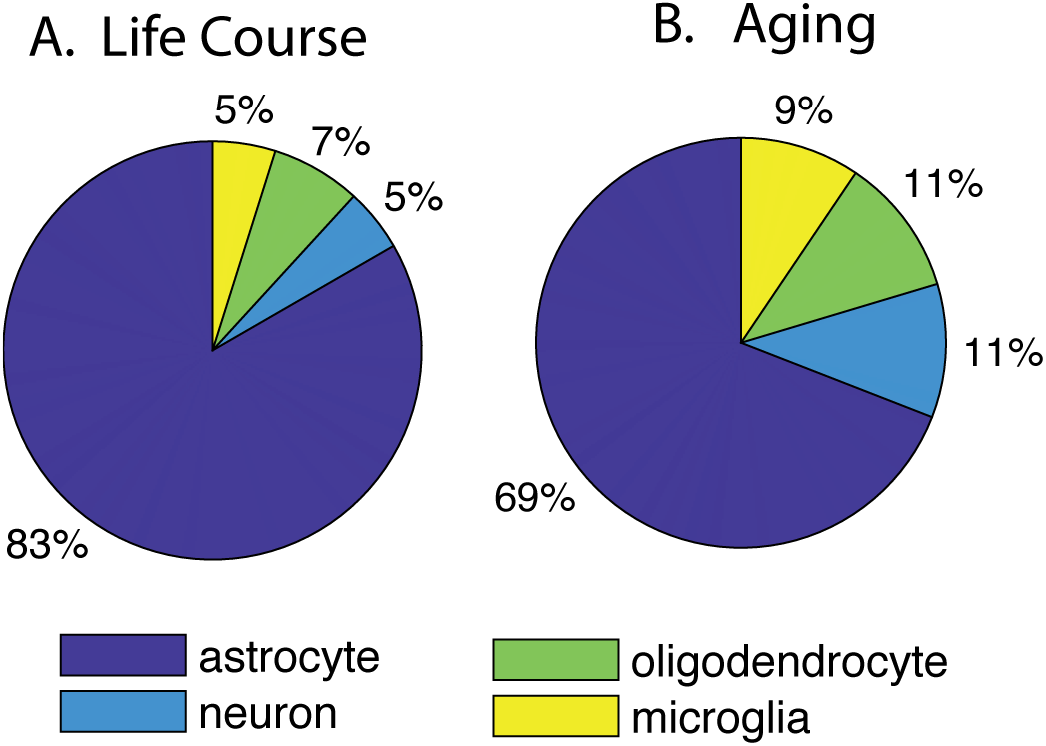
The distribution of genes in neurons, astrocytes, oligodendrocytes, and microglia in the cortex over the life course (A) and in aging (B).

**Figure 3.**
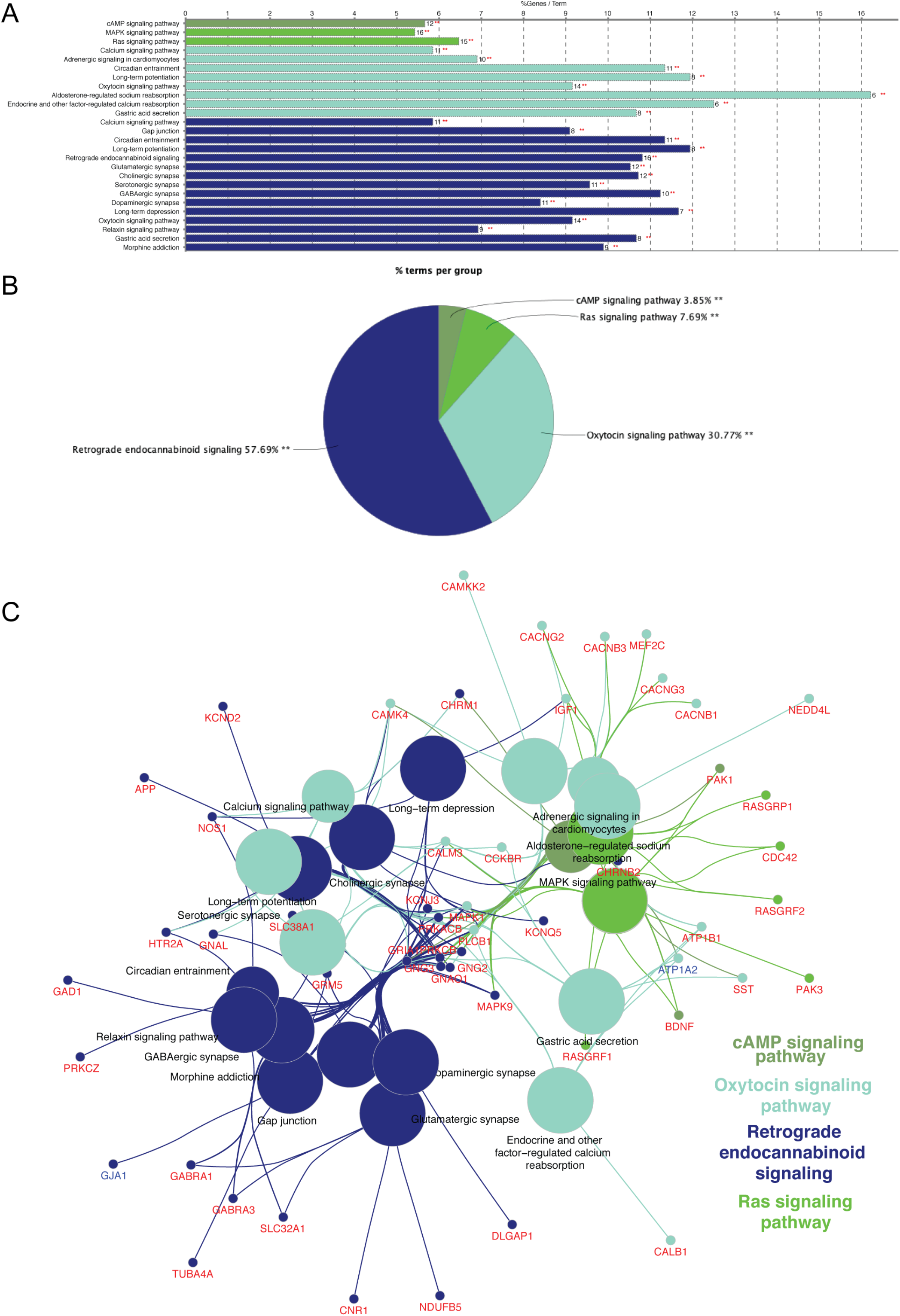
Biological pathways that were contributed by the astrocyte genes expressed in the cortex **over the life course**. Panel (A) shows individual biological pathways and the percentage of the genes contributing to each biological pathway. Panel (B) shows the four categories and the number of biological pathways in each category. Panel (C) shows the biological pathways and their related genes. The genes with the up-down and down-up expression patterns across the life course were labeled in red and blue, respectively. Each line represents the connection between the gene and biological pathway.

A hub gene was defined as having connections with 10 or more other astrocyte genes. The hub genes among the 240 astrocyte genes included GRIA1, MAPK1, APP, GRM5, BDNF, SYT1, SLC32A1, GNG3, GNG2, CDC42, SST, and CNR1, of which APP, BDNF, CDC42, SST, and CNR1 were also identified as aging-related genes published in the GenAge database (13). These genes were involved in neuroinflammation (14), the regulation of neurite growth (15), promotion of neuronal survival (16) and premature cellular senescence phenotypes (17).

### Spatio-temporal correlation between gene expression and brain morphology in aging

Next, we aimed to identify biological processes that are only relevant to aging. For this, the MRI scans of 767 subjects aged 55 years and above were extracted from the above life course dataset. The volumes of the 11 cortical regions showed significant decreases in aging (corrected *p*<0.001).

We then used linear regression to examine age effects on gene expression. Across the 11 cortical regions, 740 genes were significantly correlated with age. We applied spatio-temporal correlation analysis between gene expression and cortical volumes of the subjects aged 55 years and above to identify genes relevant to brain aging. Among them, 656 genes were correlated with cortical volumes (corrected *p* <0.05) and 282 of them were brain-dominant. As shown in **Figure 2B**, the brain-dominant genes were expressed in astrocytes (n=190), neurons (n=29), oligodendrocytes (n=30), microglia (n=26), and unclassified cells (n=2). **Table S2** lists the genes in each cell type. The neuron and microglia genes were not enriched for any biological pathway. However, the astrocyte genes were enriched for biological pathways (**Figure 4A** and **Table S4**, all *p*<0.05) that can be classified into six categories, including retrograde endocannabinoid signaling, renin-angiotensin system, proteasome, sphingolipid signaling pathway, insulin secretion, TGF-beta signaling pathway. **Figure 4C** shows the genes contributing to individual biological pathways.

**Figure 4.**
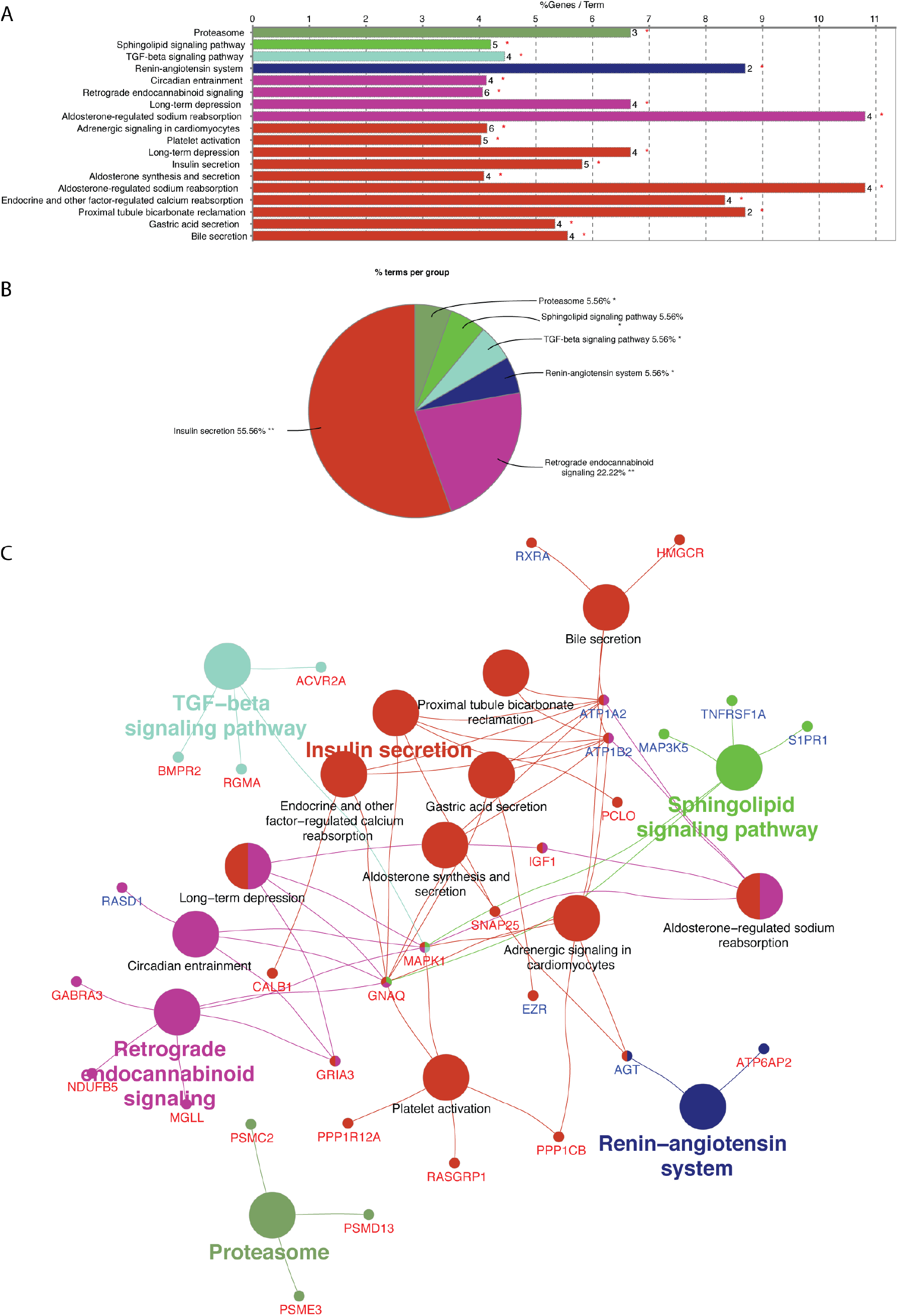
Biological pathways that were contributed by the astrocyte genes expressed in the cortex **in aging**. Panel (A) shows individual biological pathways and the percentage of the genes contributing to each biological pathway. Panel (B) shows the four categories and the number of biological pathways in each category. Panel (C) shows the biological pathways and their related genes. The genes with down-regulation and up-regulation in aging were labeled in red and blue, respectively. Each line represents the connection between the gene and biological pathway.

The hubs among the 190 astrocyte genes in the aging group included MAPK1, SNAP25, AGT, IGF1, and SYT1. Among the astrocyte genes, 61 were also discovered in the life course dataset above and were enriched for the retrograded endocannabinoid signaling (corrected p=0.0002). Moreover, 4 astrocyte genes, such as APEX1, IGF1, MAP3K5, and PCMT1, were also identified in the GenAge database (13). These genes were involved in the repair of oxidative DNA damage (18), stress response, and apoptosis (19), or the repair and degradation of damaged proteins (20).

### Trait heritability

The linkage disequilibrium score regression was used to determine whether genes involved in brain morphology across the life course and in aging were associated with 28 clinically-relevant traits related to birth outcomes (birth weight, head circumference), childhood history (preschool internalizing problems, intelligence, obesity, age at menarche), medical and socioeconomic status (HDL, LDL, BMI, cardiogram, rheumatoid arthritis, depressive symptoms, neuroticism, educational attainment, Drinking, smoking), cognition (reaction time, memory), neuropsychiatric disorders (ADHD, Autism Spectrum Disorder (ASD), schizophrenia, major depressive disorder (MDD), anxiety and bipolar disorders), neurological diseases (amyotrophic lateral sclerosis, focal and generalized epilepsy, and Alzheimer’s disease (AD)).

**Figure 5A** illustrates that the astrocyte genes in the cortex derived from the life course dataset were enriched for age at menarche, rheumatoid arthritis, depressive symptoms, neuroticism, bipolar and anxiety disorders, as well as educational attainment, drinking, and smoking. **Figure 5B** shows that the astrocyte genes in the cortex derived from the aging dataset were enriched for similar phenotypes, such as neuroticism, drinking, and smoking. But, the astrocyte genes derived from the aging dataset were also enriched for memory.

**Figure 5.**
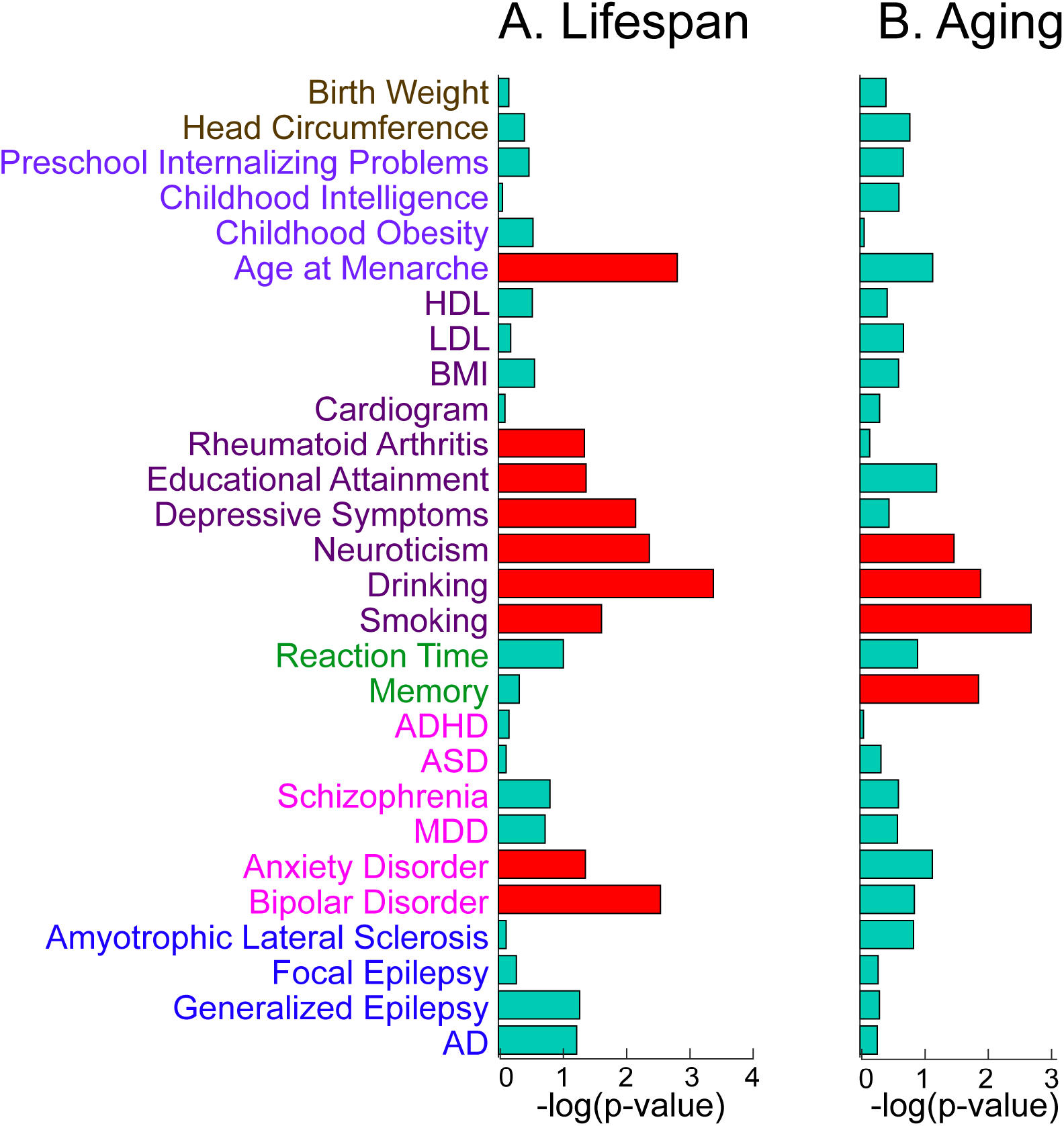
Phenotypes in associations with the astrocyte genes derived from the cortex of the life course data (A) and the aging data (B). Red bars are with statistical significance at p-values less than 0.05.

### Replication

To validate these findings, spatio-temporal correlation analysis was performed again using a replication sample of 626 subjects aged 4 to 82 years (**Section B and Table S6 in Supplementary Material**). The astrocyte genes obtained from the discovery sample significantly overlapped with those from the replication sample for the cortex (99.4%). Detailed findings were described in **Section C** of **the Supplementary Material**. The replication analysis confirmed the results reported above for the cortex.

## Discussion

This study explores the integrative analysis of neuroimaging and gene transcriptome data across the life course and provides detailed biological and phenotypical links between brain development and aging, supporting the concept of DTA. A large proportion of genes whose expression was associated with the cortical volumes across the life course were in astrocytes. These genes particularly showed up-regulation during development and the down-regulation during aging, and contributed to fundamental functions of astrocytes for maintaining homeostasis crucial for neuronal function, such as cAMP and Ras signal pathways, as well as retrograde endocannabinoid signaling. These pathways significantly overlapped with those contributed by the genes associated with cortical volume specifically during aging. Nevertheless, the genes associated with the cortical volumes in aging were also enriched for sphingolipid signaling pathway, renin-angiotensin system (RAS), proteasome, and the TGF-beta signaling pathway, which is related to the senescence-associated secretory phenotype. Both sets of genes shared common phenotypes of neuroticism, drinking, and smoking. Memory was the unique phenotype associated only with the genes relevant to cortical aging. Our findings suggest that both development and aging share common biological pathways and phenotypes. Moreover, there is a unique set of biological pathways and phenotypes associated with aging.

A previous study identified genes undergoing developmental up-regulation followed by down-regulation in aging based on the transcriptome profiles of the frontal cortex(1). Our findings revealed the presence of a gene set that was not only expressed in the frontal cortex but also the entire cortex in an up-down expression pattern across the life course. This suggests that this reversal may be a fundamental molecular trend for mirroring brain aging to brain development. Along with the findings in the transcriptome study that highlighted the neuronal genes with the up-down expression pattern across the life course (8), our study also reveals that genes associated with cortical morphology across the life course were enriched in astrocytes.

The identified genes mainly contributed to the biological pathways related to retrograde endocannabinoids, Ras, and cAMP signaling. Our study showed that the genes in these three pathways largely overlapped (**Supplementary Table S3**) with those derived from the aging dataset (**Supplementary Table S4**). For example, cAMP is one of the most common and universal second messengers in numerous signal transduction pathways. It influences the transcription of cyclin A, which promotes proliferation (21). CAMP signaling also regulates pivotal cellular functions, including cell growth and differentiation, gene transcription, and protein expression (22). Although conflicting, most studies indicate that the aging brain has a disruption in the cAMP signaling pathway, which results in reduced *cAMP response element binding protein* (CREB) activity that has been linked to long-term potential and neuroprotection (23). Whether these age-related changes are detrimental to aging or compensatory, however, remains to be resolved, as the addition of a cAMP analog exacerbates age-related memory decline, whereas blocking the pathway reverses these deficits (24–26).

Similarly, Ras proteins function as molecular switches that control intracellular signaling networks. Ras proteins coordinate multiple cellular responses, such as proliferation, differentiation, apoptosis, senescence, and metabolism. The Ras/MAPK cascade transmits signals downstream and results in the transcription of genes involved in cell growth and division (27). Activation of Ras has been considered as a factor that induces senescence (28, 29). Inhibition of the Ras/MAPK signal transduction pathway may, therefore, promote longevity by preventing cellular senescence, as has been in yeast and flies (PMID 26119340) (30, 31).

Furthermore, endocannabinoids serve as retrograde messengers at synapses in various regions of the brain. The endocannabinoids activate the CB1 receptors, such as the one encoded by the CNR1 gene, at presynaptic terminals and suppress the release of the inhibitory transmitter GABA or excitatory transmitter glutamate by inhibiting Ca2+ channels. Thus, retrograde endocannabinoid signaling is the principal mode by which endocannabinoids mediate short- and long-term forms of plasticity at both excitatory and inhibitory synapses (32). The cannabinoids have anti-inflammatory and neuroprotective actions, and the endocannabinoid activity modulates critical molecular and cellular processes influencing the aging process. Activation of cannabinoid receptors by agonist or by elevation of endocannabinoid levels promotes cell proliferation, neurogenesis, and neuronal diversification (33). Hence, the elevation of cannabinoid receptor activity either by pharmacological blockade of the degradation of cannabinoids or by receptor agonists could be a promising strategy for slowing down the progression of brain aging (33). Together, these findings suggest the existence of a set of the astrocyte genes whose expression mirrors development and aging and contributes to biological relevant pathways modulating aging.

In addition to pathways changing across the life course, our findings highlighted four unique biological pathways associated with cortical aging that are relevant to cellular senescence. The renin-angiotensin system (RAS) plays a crucial role in the regulation of renal, cardiac, and vascular physiology. Long-term RAS blockade extends the life course and prevents the age-related decline in cardiovascular (34) and metabolic function in hypertensive rats (35). The brain RAS is elevated with advanced age. Inhibition of this system may be beneficial in attenuating cognitive deficits observed in aging and neurodegenerative diseases, such as Alzheimer’s disease and post-stroke cognitive impairment (36, 37). The proteasome is a large intracellular protease whose presence is not unique in the central nervous system (38), and is responsible for the degradation of a large subset of cellular proteins (39). For the brain, the proteasome inhibition is sufficient to induce neuron death in vitro. Alterations in proteasome activity may contribute to the aging process and the elevations in protein oxidation, protein aggregation, and neurodegeneration in the aging brain (40, 41). Indeed, genetic alterations that result in enhanced proteasome function extend the life course in yeast (38). Likewise, sphingolipids are bioactive molecules that regulate cell biology and processes, such as cell cycle, senescence, proliferation, and migration (42). The bioactive sphingolipids, such as ceramide and sphingosine-1–phosphate (S1P) receptors highlighted in **Supplementary Table S4**, play crucial roles in immune responses, inflammation, cancer, metabolic and cardiovascular diseases, and neurodegeneration (43). Ceramide induces cellular senescence and S1P delays it. Alteration in the expression and in the activity of S1P metabolizing enzymes and reduced levels of S1P have been widely reported in human brains of Alzheimer’s disease (44). Administration of an S1PR selective agonist reduces spatial memory impairments and attenuates the Aβ1-induced neuronal loss in a rat model of Alzheimer’s disease (45). S1P receptors stimulate the production of neurotrophin, activate neuronal pro-survival pathways, and delay disease progression in mouse models of Huntington disease (46). Hence, the manipulation of sphingolipid metabolism may represent a great way to develop more effective and targeted therapeutic strategies. Finally, the transforming growth factor-β (TGF-β) is a superfamily of cytokines that control pleiotropic cellular functions. TGF-β is secreted as one of the senescence-associated secretory phenotype (SASP) factors and can induce and maintain the senescent phenotype, affecting cell proliferation, cell cycle regulation, DNA damage repair, telomere regulation, unfolded protein response, and autophagy (47). Aging and fibroblast senescence correlates with increased TGF-β signaling (48).

Although cellular senescence has long been linked with aging, its role in brain postnatal development has been under-investigated. Our study provided new evidence to support the potential roles of cellular senescence in brain development and aging. This idea has also been supported by the hub genes (e.g., APP, BDNF, CDC42, SST, CNR1) relevant to cellular senescence as well as the phenotypes (mood-related, smoking, drinking, age at menarche) that are relevant to longevity. Hence, our study provided the new set of the astrocyte genes and biological pathways as future targeted strategies for preventing brain aging from early brain development.

The analysis presented here combines unique imaging and transcriptome datasets. With a large sample size of brain images across the life course, we validated our results by splitting our imaging data into two independent datasets and replicating the analysis. Unfortunately, we are unable to validate our results in an independent transcriptomic dataset because there were no publicly available datasets with enough samples across the life course. Another limitation of this study is the lack of direct correspondence between the imaging and transcriptome data. A more dense temporal sampling of gene expression is needed to match imaging time points and improve the reliability of our findings. Given that we are evaluating associations across a range of ages, it is possible that the significant associations observed are relevant for cortical structure throughout the life course.

In summary, we demonstrate that cortical morphology is significantly associated with the expression profiles of astrocyte markers across the life course. These associations reflect shared biological pathways of the cortical development and aging. These findings suggest the genes and biological mechanisms mirroring development and aging that can be potential targets for preventing aging from development.

## Methods

### Gene expression dataset and analysis

Kang’s et al (2011) transcriptome dataset (11) was retrieved through the NCBI Gene Expression Omnibus Accession number GSE25219. Affymetrix GeneChip Human Exon 1.0 ST Array platform was used to measure the expression level of genes at 11 cortical regions in each donor’s brain. **Supplementary Table S1** lists the 11 cortical regions sampled in Kang’s study (11). Partek Genomics Suite version 6.5 (Partek Incorporated, St. Louis, MO, USA) was employed to perform robust multi-array analysis (RMA) background correction, quantile normalization and log2. Each probe was annotated with gene symbol obtained from the GPL file deposited in GEO, leading to a total of 17565 genes to be investigated in this study. This study included postnatal post-mortem brains of 26 donors aged 2 to 82 years, corresponding to the age range of the neuroimaging sample. As the number of donors being sampled in each hemisphere and age point were different, gene expression values were averaged across hemispheres and donors within the 13 available age points: Age 3 (2-3 years), Age 4 (4 years), Age 8 (8 years), Age 15 (11-15 years), Age 19 (18-19 years), Age 20 (21-23 years), Age 30 (30-37 years), Age 40 (40 years), Age 50 (55 years), Age 60 (64 years), Age 70 (70 years) and Age 80 (82 years).

### T1-weighted MRI datasets and analysis

T1-weighted MRI data of 2765 healthy subjects aged 3 to 82-years-old from six were obtained from publicly available datasets, including Pediatric Imaging, Neurocognition, and Genetics (PING)(49); Attention Deficit Hyperactivity Disorder 200 (ADHD-200)(50); Nathan Kline Institute-Rockland Sample (51, 52); 1000 Functional Connectomes Project data (FCP)(53) and Southwest University Adult Lifespan Dataset (SALD)(54). The demographic information of each dataset is described in **Supplementary Table 5**. The exclusion criteria were existing diagnosis of chronic medical conditions (e.g., cancer, congenital abnormalities), mental illnesses (e.g., ADHD, Autism), neurological disorders and/or head injury with loss of consciousness.

The structural MRI datasets were analyzed using FreeSurfer 5.3 (http://surfer.nmr.mgh.harvard.edu). Cortical volumes corresponding to the aforementioned regions listed in **Supplementary Table S1** were extracted.

### Statistical analysis

Age (in years) was log2 transformed because of its proper representation for phenotypes across the life course (55, 56). We examined 1) the effects of age on gene expression levels and brain volumes across the life course; 2) spatio-temporal correlation between gene expression and brain volume across the life course; 3) repeated analysis from 1) and 2) using only the datasets with age above 55 years.

A quadratic regression model was used to test age effects on the expression profile of each gene in every brain region (57–59). The model included linear and quadratic terms of age entered as main factors and the expression level of each gene entered as the dependent variable (expression level of each gene in each brain region ~ β_0_ + β_1_ Age + β_2_ Age^2^ + *ε*). The genes whose expression is as a quadratic function of age in the same direction within all the 11 cortical regions, i.e., an up-down pattern or down-up pattern across the life course, were identified.

The same quadratic analysis was applied to the cortical volume data. We identified quadratic age effects on the volume of the 11 cortical regions across the life course (brain volume ~ β_0_ + β_1_ Age + β_2_ Age^2^ + *ε*). Statistical p-values (p<0.05) were corrected using False Discovery Rate (FDR).

Pearson’s correlation analysis was used to examine spatio-temporal associations between gene expression and cortical morphology across the life course. The cortical volumes were averaged across hemispheres and individuals within the 13 age groups. Pearson’s correlation analysis was used to investigate the association between each gene’s expression value (expression values from 11 cortical regions × 13 age groups) and brain volumes (11 cortical volumes × 13 age group). FDR correction was used to correct for multiple comparisons over the genes at p<0.05.

Finally, we limited our analysis to the datasets with age above 55 years. The model included a linear term of age entered as a main factor and the expression level of each gene entered as the dependent variable (expression level of each gene in each brain region ~ β_0_ + β_1_ Age + *ε*) with FDR correction (p<0.05). Genes that showed expression in an up pattern or down pattern in aging across all the cortical regions were identified.

The same linear analysis was applied to the cortical volume in the aging dataset with FDR correction for multiple comparisons over the cortical regions (p<0.05). The above spatio-temporal correlation between gene expression and cortical morphology in aging was also carried out using FDR correction of multiple comparisons over the genes (p<0.05).

### Gene functional analysis

#### Brain-specific genes

This study employed TISSUES 2.0 (https://tissues.jensenlab.org) to identify brain-specific genes. TISSUES 2.0 provides a confidence score on the association between protein-coding genes and tissues based on transcriptomics, proteomics data that were manually curated and automatically text-mined information from biomedical literature. Confidence scores can range from 0 to 5. A higher score indicates greater confidence that a gene is expressed or present in a certain tissue. This study identified the genes with the confidence score associated with brain tissue greater than 4.5.

#### Cell types

We based on the cortical transcriptome database (http://www.brainrnaseq.org) (60) to assign each gene a cell type, including neuron, astrocyte, microglia, oligodendrocyte. This database contains the expression level of 23222 genes in these four cell types. For each gene, the cell type with the highest expression level was annotated as the preferentially expressed cell type.

#### Biological pathways and Gene hub analysis

Biological pathways of cell-specific genes were analyzed based on KEGG using Cytoscape ClueGo (61). Hypergeometric distribution tests were performed to identify enriched biological pathways (FDR corrected p<0.05). ClueGo grouped biological pathways based on the presence of genes among them. This study illustrated the findings of individual biological pathways as well as their groups.

We employed web-based STRING (62) to identify protein-protein interactions that were derived from high-throughput experiments, literature mining, gene fusion, co-occurrence, coexpression analysis, and genomic meta-analysis. The connections among the genes were formed through this analysis. The degree of each gene was defined as the number of its connections with other genes.

### Linkage disequilibrium score regression

Linkage disequilibrium (LD) score regression (https://github.com/bulik/ldsc/) was used to evaluate the trait heritability of a gene set. An LD score regression employs Genome-Wise Association Study (63) summary statistics and regresses the association test statistics of SNPs on their LD scores. This study obtained GWAS summary statistics from Genome-Wide Repository of Associations between SNPs and traits (https://grasp.nhlbi.nih.gov/). The trait summary statistics of 29 GWAS studies with a sample size greater than 5000 were included in this study. Genetic variants of each gene were extracted within ±10kb using R. Annotation-specific LD scores were further estimated using 1000 Genomes samples with European ancestry, and a default 1-centiMorgan window (64). The LD score of an SNP is the sum of LD *r*^2^ measured with all other SNPs. Finally, for each trait, LDSC regresses the χ^2^ (or Z^2^) association statistic from each SNP onto the “LD-score” of that SNP to evaluate the contribution of the genome annotation to trait heritability. Annotation categories with a significant positive regression coefficient denoted that the annotation positively contributed to the trait with above-average heritability, while negative regression coefficient denoted that the annotation negatively contributed to the trait with below-average heritability(65). Similar to previous studies (64, 65), p-values for the traits with a significant positive regression coefficient were reported in this study.

#### Data-accessibility

Please contact the corresponding author regarding access to data used in this paper.

## Acknowledgments

This research is supported by the Singapore National Research Foundation under its Translational and Clinical Research (TCR) Flagship Programme and administered by the Singapore Ministry of Health’s National Medical Research Council (NMRC), Singapore-NMRC/TCR/004-NUS/2008; NMRC/TCR/012-NUHS/2014. Additional funding is provided by the Singapore Ministry of Education (Academic research fund tier 1; NUHSRO/2017/052/T1-SRP-Partnership/01), and NUS Institute of Data Science, Singapore.

## Supplementary Material

**Supplementary Table S1.**
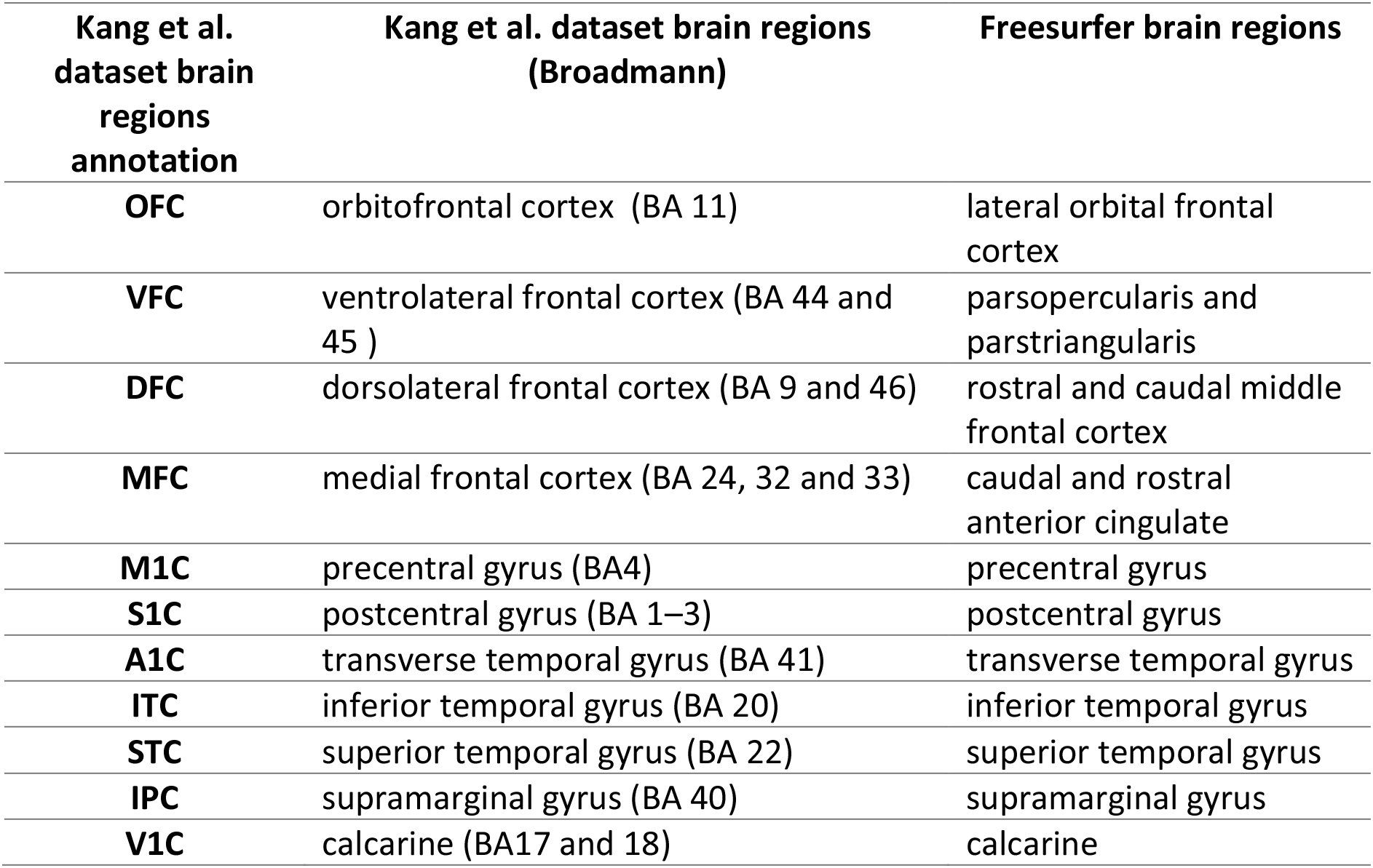
Transcriptome and Freesurfer brain anatomical regions.

**Supplementary Figure S1.**
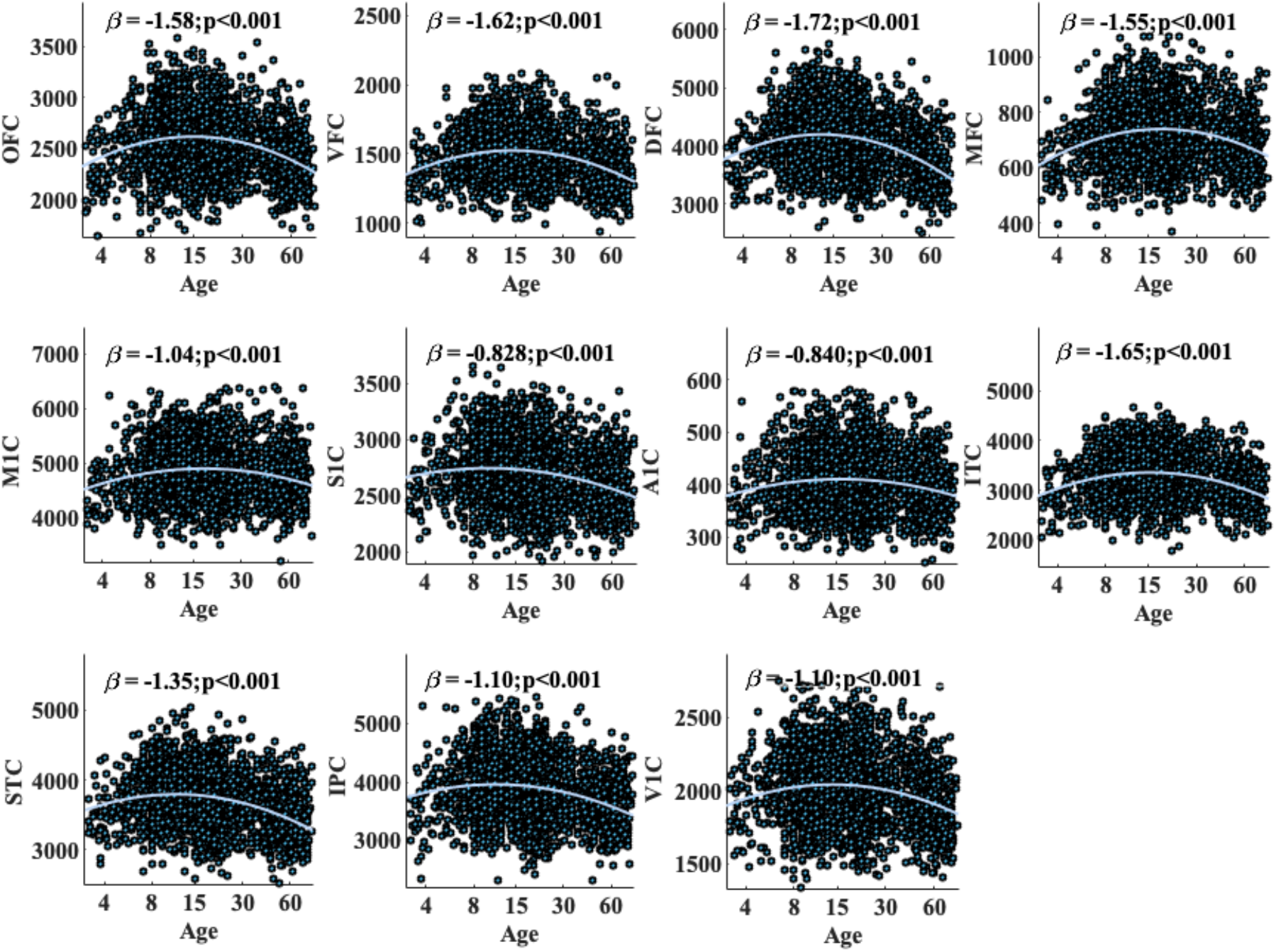
Quadratic age effects on cortical volumes in the discovery dataset over the life course. Abbreviations: OFC, orbitofrontal cortex; VFC, ventrolateral prefrontal cortex; DFC, dorsolateral prefrontal cortex; MFC, medial frontal cortex; M1C, motor cortex; S1C, somatosensory cortex; A1C, transverse temporal cortex; ITC, inferior temporal cortex; STC, superior temporal cortex; IPC, inferior parietal cortex; V1C, calcarine.

**Supplementary Table S2.**
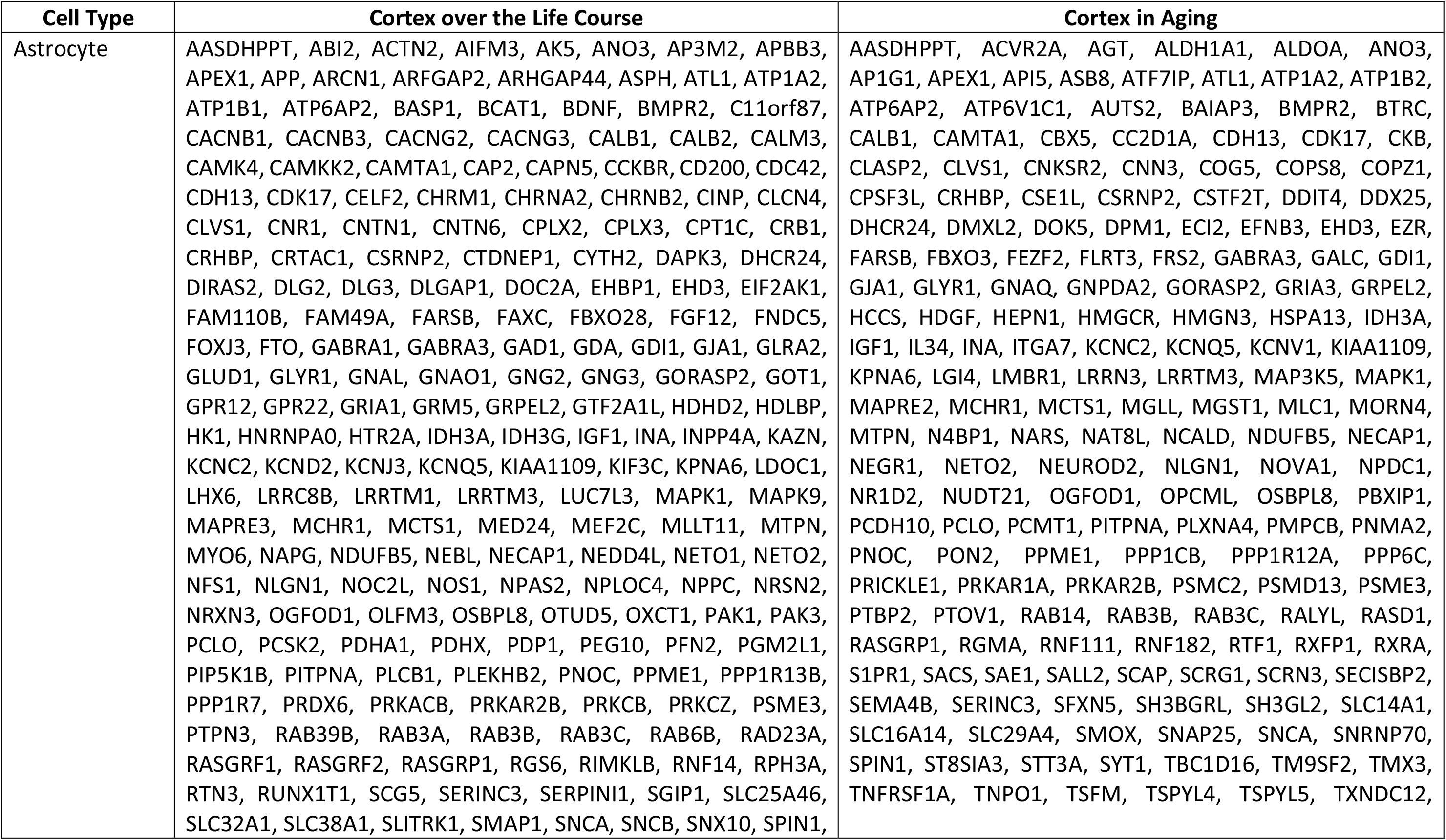
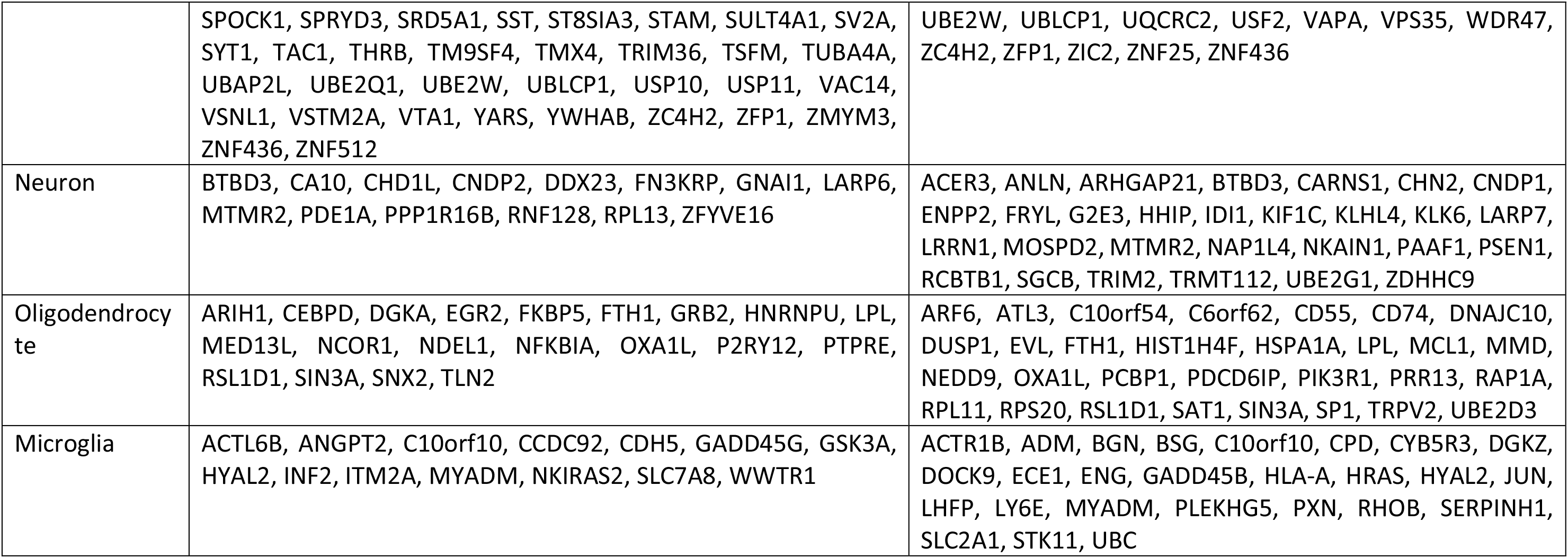
The list of genes whose expression pattern was significantly correlated with the cortical morphology in the life course (2^nd^ column) and aging datasets (3^rd^ column).

**Supplementary Table S3.**
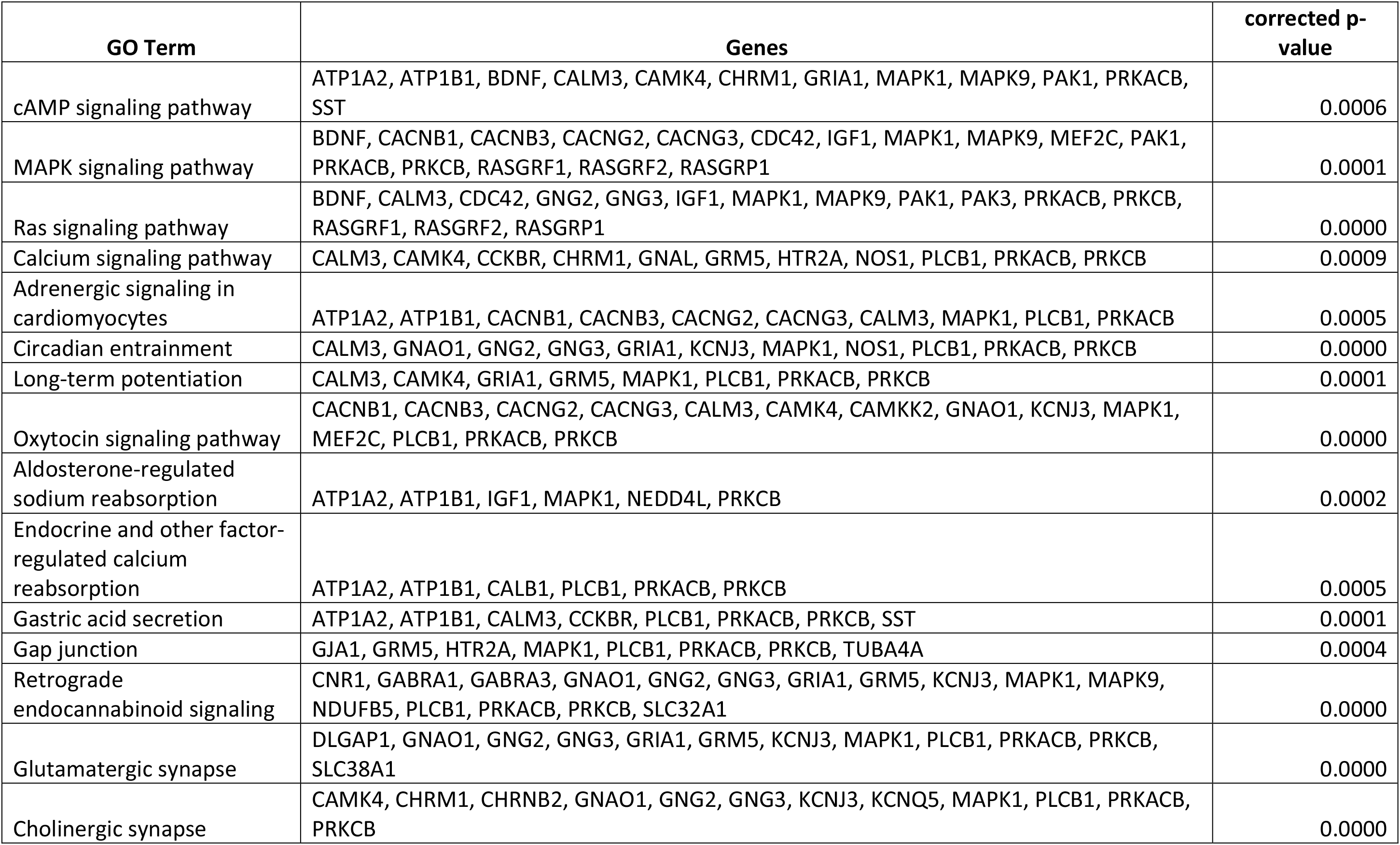
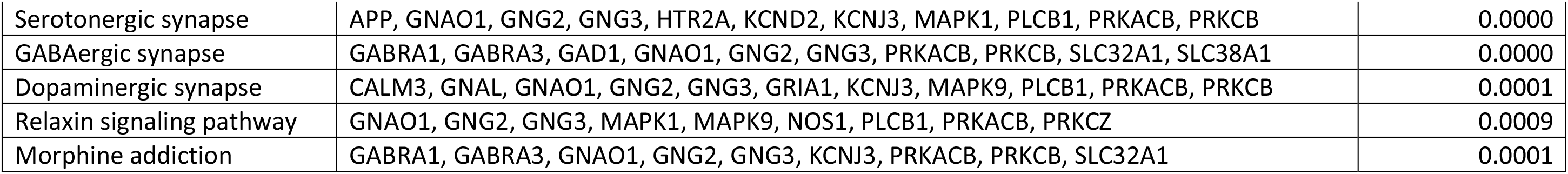
Biological pathways contributed by the genes whose expression pattern was associated with the cortical morphology over the life course.

**Supplementary Table S4.** Biological pathways contributed by the genes whose expression pattern was associated with cortical morphology in aging.

| GO Term | Genes | corrected p-values |
| --- | --- | --- |
| Proteasome | PSMC2, PSMD13, PSME3 | 0.026 |
| Sphingolipid signaling pathway | GNAQ, MAP3K5, MAPK1, S1PR1, TNFRSF1A | 0.024 |
| TGF-beta signaling pathway | ACVR2A, BMPR2, MAPK1, RGMA | 0.032 |
| Renin-angiotensin system | AGT, ATP6AP2 | 0.039 |
| Circadian entrainment | GNAQ, GRIA3, MAPK1, RASD1 | 0.037 |
| Retrograde endocannabinoid signaling | GABRA3, GNAQ, GRIA3, MAPK1, MGLL, NDUFB5 | 0.021 |
| Long-term depression | GNAQ, GRIA3, IGF1, MAPK1 | 0.023 |
| Aldosterone-regulated sodium reabsorption | ATP1A2, ATP1B2, IGF1, MAPK1 | 0.015 |
| Adrenergic signaling in cardiomyocytes | AGT, ATP1A2, ATP1B2, GNAQ, MAPK1, PPP1CB | 0.023 |
| Platelet activation | GNAQ, MAPK1, PPP1CB, PPP1R12A, RASGRP1 | 0.026 |
| Insulin secretion | ATP1A2, ATP1B2, GNAQ, PCLO, SNAP25 | 0.018 |
| Aldosterone synthesis and secretion | AGT, ATP1A2, ATP1B2, GNAQ | 0.036 |
| Endocrine and other factor-regulated calcium reabsorption | ATP1A2, ATP1B2, CALB1, GNAQ | 0.020 |
| Proximal tubule bicarbonate reclamation | ATP1A2, ATP1B2 | 0.039 |
| Gastric acid secretion | ATP1A2, ATP1B2, EZR, GNAQ | 0.026 |
| Bile secretion | ATP1A2, ATP1B2, HMGCR, RXRA | 0.025 |

## Section A. Discovery Sample

**Supplementary Table S5** lists the demographic data of 2765 subjects with T1 structural MRI in the discovery sample for our main analysis. Details on the source of each T1 structural MRI dataset were described here.

**Supplementary Table S5.** Demographics of 2765 subjects with structural MRI data.

| Dataset | N | Age Range | Age mean (SD) |
| --- | --- | --- | --- |
| PING | 809 | 3-21 | 12.3 (5.06) |
| ADHD200 | 548 | 7-26 | 11.7 (3.53) |
| 1000 FCP | 733 | 8-79 | 27.7 (13.5) |
| SALD | 494 | 19-80 | 45.2 (17.4) |
| NKI | 181 | 6-82 | 36.7 (20.0) |

Abbreviations: PING, Paediatric Imaging, Neurocognition, and Genetics; ADHD, Attention Deficit Hyperactivity Disorder 200; FCP, 1000 Functional Connectomes Project; SALD, Southwest University Adult Lifespan Dataset; NKI, Nathan Kline Institute-Rockland Sample.

**PING.** The PING applied a standardized high-resolution MRI protocol, including T1-weighted scans, across the nine sites and 12 scanners. The PING T1 MRI protocol for each scanner manufacturer is provided on http://pingstudy.ucsd.edu/resources/neuroimaging-cores.html.

**ADHD200.** Data for ADHD-200 was collected as a large collaborative effort across 8 sites. T1-weighted volumetric Magnetization Prepared Rapid Gradient Echo (MPRAGE) structural MRI was acquired on every subject and MRI protocol for each site is provided on http://fcon_1000.projects.nitrc.org/indi/adhd200/.

**1000 FCP.** A high-resolution T1-weighted anatomical image was obtained for each subject. The general data acquisition procedures were similar across 7 sites and MRI protocol for each site is provided on http://fcon_1000.projects.nitrc.org.

**SALD.** All adults were scanned using a 3.0 T Siemens Trio MRI scanner at the Southwest University Center for Brain Imaging. The image protocol was high-resolution T1-weighted anatomical images prepared rapid gradient echo (MPRAGE; 176 slices, 1 mm thickness, matrix 256×256, repetition time 1900 ms, echo time 2.52 ms, flip angle 90°).

**Nathan Kline Institute-Rockland Sample (NKI)**. All subjects were scanned using a 3T Siemens TIM Trio system. The image protocol was high-resolution T1-weighted anatomical images prepared rapid gradient echo (MPRAGE; 176 slices, 1 mm thickness, repetition time 1900 ms, echo time 2.52 ms, flip angle 9°). Data is obtainable from http://fcon_1000.projects.nitrc.org/indi/enhanced/.

## Section B. Replication Study

### Replication Sample

A total of 626 healthy subjects aged 4 to 82-years-old from Autism Brain Imaging Data Exchange dataset (ABIDE), local children cohort from the Cognition and Brain Development in Children (CBDC) project (Zhong et al. 2014) and local aging cohorts from the Brain and Cognition Aging Study (BCAS) (Lee et al. 2013, 2015) were leveraged to provide a normative replication sample across the life course. Both local cohort studies were approved by the National University of Singapore Institutional Review Board and all participants provided written informed consent prior to participation. In general, we excluded subjects with an existing diagnosis of chronic medical conditions (e.g., cancer, congenital abnormalities), mental illnesses (e.g., ADHD, Autism), neurological disorders and/or head injury with loss of consciousness. **Supplementary Table 6** listed the demographics of subjects included in this study.

**Supplementary Table S6.** Demographics of 626 subjects with structural MRI data for the replication sample.

| Replication Sample |  |  |  |  |  |
| --- | --- | --- | --- | --- | --- |
| Type | Database | N | Age Range | Age mean (SD) | Female % |
| Public | ABIDE | 186 | 6-56 | 16.9 (7.67) | 17.3 |
| Local | CBDC | 100 | 4-10 | 5.91 (0.81) | 52.6 |
| Local | BCAS | 340 | 22-80 | 63.5 (14.4) | 51.0 |

Abbreviations: ABIDE, Autism Brain Imaging Data Exchange; CBDC, the Cognition and Brain Development in Children project; BCAS, the Brain and Cognition Aging Study; SD, standard deviation.

**ABIDE.** T1-weighted volumetric Magnetization Prepared Rapid Gradient Echo (MPRAGE) structural MRI was acquired on every subject in the ABIDE (DiMartino et al., 2013), and the parameters used at each site can be found at http://fcon_1000.projects.nitric.org/indi/abide/.

**CBDC.** Children underwent MRI scans using a 3T Siemens Skyra scanner with a 32-channel head coil at Clinical Imaging Research Centre of the National University of Singapore. The imaging protocol was high-resolution isotropic T1-weighted Magnetization Prepared Rapid Gradient Recalled Echo (MPRAGE; 190 slices, 1mm thickness, in-plane resolution 1 mm, sagittal acquisition, field-of-view 190×190 mm, matrix=190×190, repetition time=2000 ms, echo time=2.08 ms, inversion time=850 ms, flip angle=9°).

**BCAS.** All adults were scanned using a 3T Siemens Magnetom Trio Tim scanner with a 32-channel head coil at the Clinical Imaging Research Centre of the National University of Singapore. The image protocol was high-resolution isotropic T1-weighted Magnetization Prepared Rapid Gradient Recalled Echo (MPRAGE; 192 slices, 1 mm thickness, sagittal acquisition, field of view 256×256 mm, matrix 256×256, repetition time 2300 ms, echo time 1.90 ms, inversion time 900 ms, flip angle 9°).

### Structural MRI Analysis

MRI processing for all datasets were performed with the FreeSurfer 5.3 image analysis suite (http://surfer.nmr.mgh.harvard.edu). Eleven cortical and four subcortical brain volumes corresponding to the aforementioned brain regions listed in Kang et al (2011)’s were extracted (see **Supplementary Table 1**).

### Cortical morphological pattern over the life course

The brain volumes for cortical, amygdala-hippocampal complex, and subcortical regions were estimated using FreeSurfer from the T1-weighted MRI of 626 subjects aged 4 to 82-years-old. Quadratic linear regression analysis revealed all 11 cortical volumes as a function of age^2^, suggesting a significant inverted-U relationship with age over the life course in all these brain regions (corrected *p*<0.001).

**Supplementary Figure S3.**
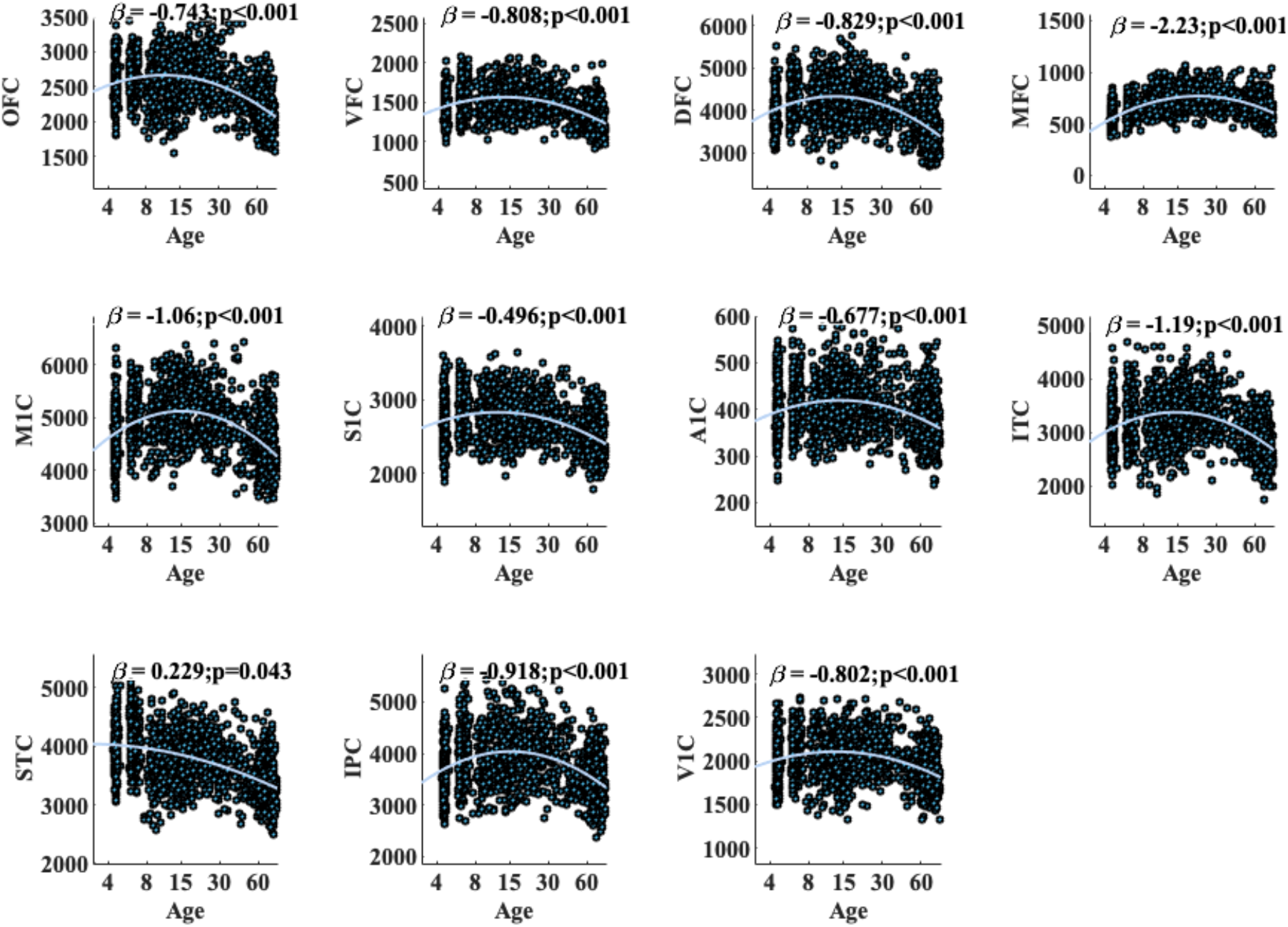
Quadratic age effects on cortical volumes in the replication sample. Abbreviations: OFC, orbitofrontal cortex; VFC, ventrolateral prefrontal cortex; DFC, dorsolateral prefrontal cortex; MFC, medial frontal cortex; M1C, motor cortex; S1C, somatosensory cortex; A1C, transverse temporal cortex; ITC, inferior temporal cortex; STC, superior temporal cortex; IPC, inferior parietal cortex; V1C, calcarine.

### Spatio-temporal correlation between gene expression and cortical morphology over the life course

Among the 710 genes with U-shaped or inverted-U gene expression profiles across life course in the cortex, 452 genes showed significant spatio-temporal correlation with the cortical volumes (corrected *p*<0.05). Among them, 227 genes were brain-dominant, which consisted of 180 astrocytes, 12 neurons, 19 oligodendrocytes, 12 microglia, and 4 unclassified cells. Among 180 astrocytes, 179 were overlapped with those revealed in the discovery sample.

## Notes

### Competing Interest Statement

The authors have declared no competing interest.

